# Refined shoulder kinematics via markerless bony landmark detection and acromial 3D shape using an RGB-D camera during hand-cycling

**DOI:** 10.64898/2025.12.04.692368

**Authors:** Amedeo Ceglia, Lucas Mulhaupt, Florent Moissenet, Mickël Begon, Lama Seoud

## Abstract

Biomechanical biofeedback has the potential to enhance rehabilitation by providing clinicians with objective evaluation of patient performances. As feedback systems often depend on expensive and sophisticated motion capture technologies, researchers explore computer vision-based alternatives. Existing methods suffer from substantial joint angle errors, particularly in the upper limb, and neglect the scapular movements. We developed an approach for detecting bony landmarks and performing refined upper-limb kinematics assessments using a single consumer-grade depth-sensing camera. Unlike other markerless methods, our model incorporates the scapula, offering comprehensive shoulder joint kinematics. Annotated images from eight participants were used to fine-tune a convolutional neural network, which was subsequently evaluated on a hand-cycling motion. Our method showed a strong agreement with a reference marker-based system, with 3D bony landmark detection errors averaging 5 mm. The resulting kinematics closely aligned with the reference system, maintaining acceptable joint angle errors (∼6.3°). Furthermore, the algorithm could provide real-time bony landmark positions and joint kinematics at a rate of 50 Hz. This study highlights the potential of using a single consumer-grade depth-sensing camera combined with an acromial 3D-shape to accurately estimate upper-limb kinematics through bony landmark detection, paving the way for more accessible clinical assessments.

## 1. Introduction

Human motion analysis is a fundamental and effective tool used in diagnosing, treating, and preventing musculoskeletal disorders. It can be useful in physical rehabilitation to provide real-time feedback and to quantify abnormal motion patterns. Objective quantification of upper limb kinematics during functional tasks is an important step toward investigating upper-limb movement disorders and assessing the effect of rehabilitation interventions (Prochazka and Kowalczewski, 2015; Kanzler et al., 2020). However, analyzing the upper-limb kinematics can be challenging due to the difficulty in tracking bones, such as the scapula and the clavicle, which slide under the skin. Upper-limb motion can be captured using various systems, including optical marker-based, inertial, or optical markerless systems.

Optical marker-based technology uses expensive optoelectronic cameras to track the 3D position of active or passive markers placed on bony landmarks. It is accurate, with marker 3D position errors of less than a millimeter (Topley and Richards, 2020), making it suitable for clinical applications. This technology is often used as a reference for assessing emerging motion capture systems. Yet, placing skin markers can be time-consuming and requires participants to wear tight or minimal clothing during data collection. Furthermore, optoelectronic systems’ high cost and setup complexity pose challenges in clinical environments (Mitchell et al., 2023).

Inertial motion capture involves placing IMU on the limb to record its orientation over time (Niswander et al., 2020). IMU has the advantages of portability, ease of use, and compatibility with regular clothing. However, gyroscopes may experience drift during long tasks, and electromagnetic fields can affect the magnetometer, leading to errors in joint angle estimates. Errors of about 10° have been reported for the lower limb (Shuai et al., 2022) and up to 30° for the upper limb (Henschke et al., 2022) when compared to marker-based systems. Another limitation is the oversimplification of the upper-limb model, which often uses a single humerothoracic joint instead of modeling the sternoclavicular, acromioclavicular, and glenohumeral joints. This is due to the complexity of capturing scapula bone under the skin (Illyés and Kiss, 2006; van Andel et al., 2009). Consequently, the role of the scapula, which is significant in upper limb rehabilitation (Voight and Thomson, 2000), is omitted.

Optical-based markerless systems are based on feature extraction from images using computer vision and machine learning algorithms. To provide 3D positioning, these systems typically use either multiple calibrated color (RGB) cameras (Lahkar et al., 2022) or a single consumer-grade depth-sensing camera (RGB-D) (Cai et al., 2019). The former offers high accuracy, when calibration is well performed, but is expensive for clinical use due to the synchronized high frequency and resolution of the cameras and required large space for installation. On the other hand, RGB-D cameras provide acceptable spatial accuracy without calibration (already performed by the manufacturer), are easy to use and cost-effective, but have a limited field of view. While RGB-D cameras may be suitable for clinical environments (Boldo et al., 2024), the existing methods may lack accuracy when applied to the upper limb. Errors in joint angle estimation can be as high as 23° (Lahkar et al., 2022). The accuracy also depends on the quality of the annotated images used to train the models. As these annotated images are often not publicly available, assessing the training quality of these models is challenging. Moreover, due to the difficulty in visualizing bones like the scapula and clavicle in images, these methods often ignore them and rely on the same oversimplifications of the shoulder complex as IMU systems. Consequently, scapular and clavicular motions are excluded, potentially resulting in an incomplete assessment of upper limb motion. While markerless systems may suit clinical environments, the omission of scapular representation limits their clinical applicability. Scapular orientation can be estimated, with a marker-based system, using an acromial marker cluster suitable for dynamic motion (Karduna et al., 2001; van Andel et al., 2009; MacLean et al., 2014). However, acromial clusters require optical calibration and are incompatible with markerless systems.

In this study, our objective was to achieve pose estimation using a single RGB-D camera by detecting key points in the depth images using a trained convolutional neural network (CNN). CNNs are commonly employed for such tasks (Jogin et al., 2018). Mathis et al. (2018) introduced the DeepLabCut (DLC) software, which provides an intuitive interface for fine-tuning, evaluating, and running numerous CNN models inference in live (Kane et al., 2020). Additionally, we aimed to capture scapular motion through RGB-D imaging, addressing a critical gap in upper-limb motion analysis. To this end, this study proposes a novel markerless approach by accurately locating bony landmarks from a single RGB-D camera to estimate the kinematics of the shoulder complex. Additionally, we introduced a custom-made acromial shape that allows the capture of the scapula from the RGB-D camera without the need for optical calibration. Therefore, our kinematic model includes both the scapula and clavicle to offer more refined upper-limb kinematics reconstruction. Additionally, we proposed a method for automatically and efficiently generating a new set of annotated images from experimental data, combined with a model fine-tuning process.

## 2. Method

This section outlines the complete process for extracting kinematics from RGB-D images. It covers data generation (Sec. 2.1), CNN model fine-tuning for the motion of interest (Sec. 2.2-2.3), and kinematics estimation based on bony landmark detection combined with the acromial 3D shape (Sec. 2.4). Our approach was evaluated using hand-cycling motion, a representative task of rehabilitation activities. Evaluation (Sec. 2.5) was performed by comparing the kinematic data obtained from both the proposed and reference motion capture systems, with a focus on the accuracy of bony landmark detection and joint kinematics.

### 2.1. Data collection and annotation

This experiment was approved by the ethics committee of the Université de Montréal (#2023-4743). Eight healthy adults (3 females; age: 25.9 ± 12.2 years old; height: 175.8 ± 8.5 cm; mass: 70.2 ± 8.4 kg) participated in the study. Thirteen squared white markers (∼2×2 cm) were placed on the trunk and left upper limb at predefined locations to locate key bony landmarks and reduce soft tissue artifacts (Blache et al., 2017). Additionally, a custom-made acromial 3D shape, with 3 white markers, was placed on the participants to estimate the scapula’s bony landmarks (acromial angle, trignium spinae, and inferior angle). This shape enabled the estimation of the scapula’s orientation from a frontal perspective without requiring optical calibration, with the same level of accuracy as acromial clusters presented in the literature (see Appendix A).

Participants were asked to perform 2 min-hand pedal trials on a stationary hand-cycle at 15, 20, 30, and 40 W all performed at 60 revolutions per minute. A single stereo-vision RGB-D camera (RealSense D455, Intel, USA) was used to record both color (RGB) and depth images at 60 Hz with a 848×480 pixels resolution each. It was placed approximately one meter away from the thorax, roughly aligned with the participant’s neck (Fig. 2). For the validation, a 14-camera motion capture system (T40s, Vicon, Oxford, UK) was used to record (120 Hz) 13 hemispherical (ø 1 mm) retro-reflective markers placed in the center of the white markers.

Part of the method to automatically retrieve the bony landmarks (Sec. 2.4.1) relied on automatic key point detection in the depth images using a CNN. For training purposes, the key points’ positions were annotated on the depth-aligned RGB images, following the approach described in Ceglia et al. (2025). In brief, a tracking algorithm was developed that involves manually labeling the first frame, followed by applying tracking algorithms to follow each marker’s trajectory based on color (optical flow) and velocity (Kalman filter). The use of the 3D space provided by the camera further enhances the accuracy of the tracking. To generate the training dataset, we used this method on a 2-minute trial, after removing all reflective markers. Thanks to the automatic labeling method proposed in Ceglia et al. (2025), we could provide accurate (*<* 4 mm error from reference system) and fast annotations (around 60 FPS) for more than 6 000 depth images per participant.

### 2.2. Image preprocessing

A first preprocessing step consisted in background removal on all recordings using an empirical threshold of 1.2 m on the depth image to eliminate all associated artifacts. The images were also cropped around the participant to reduce the size and thus the subsequent processing time. The algorithm aims to automatically detect bony landmarks on the preprocessed depth images to allow for kinematics estimation. Therefore, to leverage existing pre-trained CNN architectures, the pre-processed depth images (single-channel, 16-bit format) were converted into color-like images with three channels in 8-bit format. This transformation was achieved by computing the surface normals, as proposed by Wang and Siddiqi (2016). The partial derivative method (Monga and Benayoun, 1995) was employed to ensure fast computation of the surface normals. This approach had two main objectives: first, to enable the use of the pre-trained model provided by DLC, and second, to generate data that reflect the orientation of the surface, which can be beneficial for detecting key features on the skin surface. The surface was defined by the function:

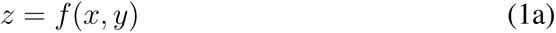

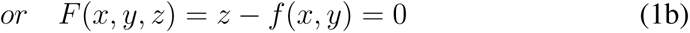

where *x* and *y* are the pixel location along the width and height dimensions, and *z* is the depth value. The normal to the surface (*n*) is equal to the gradient of the function (∇*F* (*x, y, z*)):

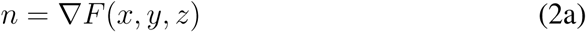

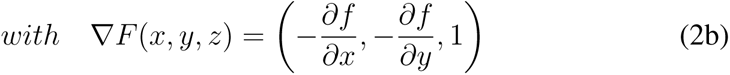

The normal vector was then normalized and mapped to the range [0, 255]. The transformed 3-channels depth image was built using the normalized vector components, with *x*, *y*, and *z* being assigned to the R, G, and B channels respectively (Fig. 1). To improve the generalization of the model on altered-quality images, one-third of the training images was downsampled with a homogeneous ratio of 0.8, another third with a ratio of 0.9, and the last third was left unchanged.

**Figure 1:**
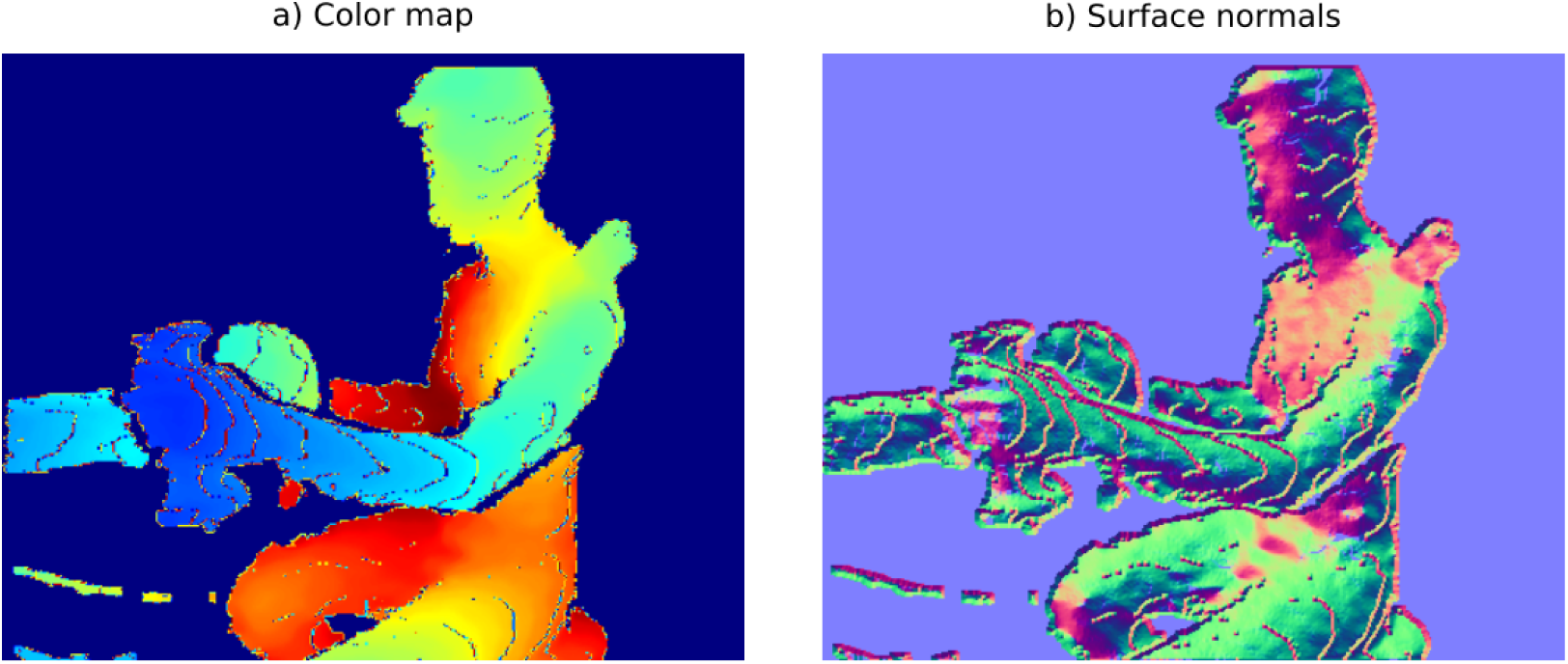
Cropped depth image, for one participant, after removing the background, and a) applying a color map or b) computing the surface normals from the depth image.

**Figure 2:**
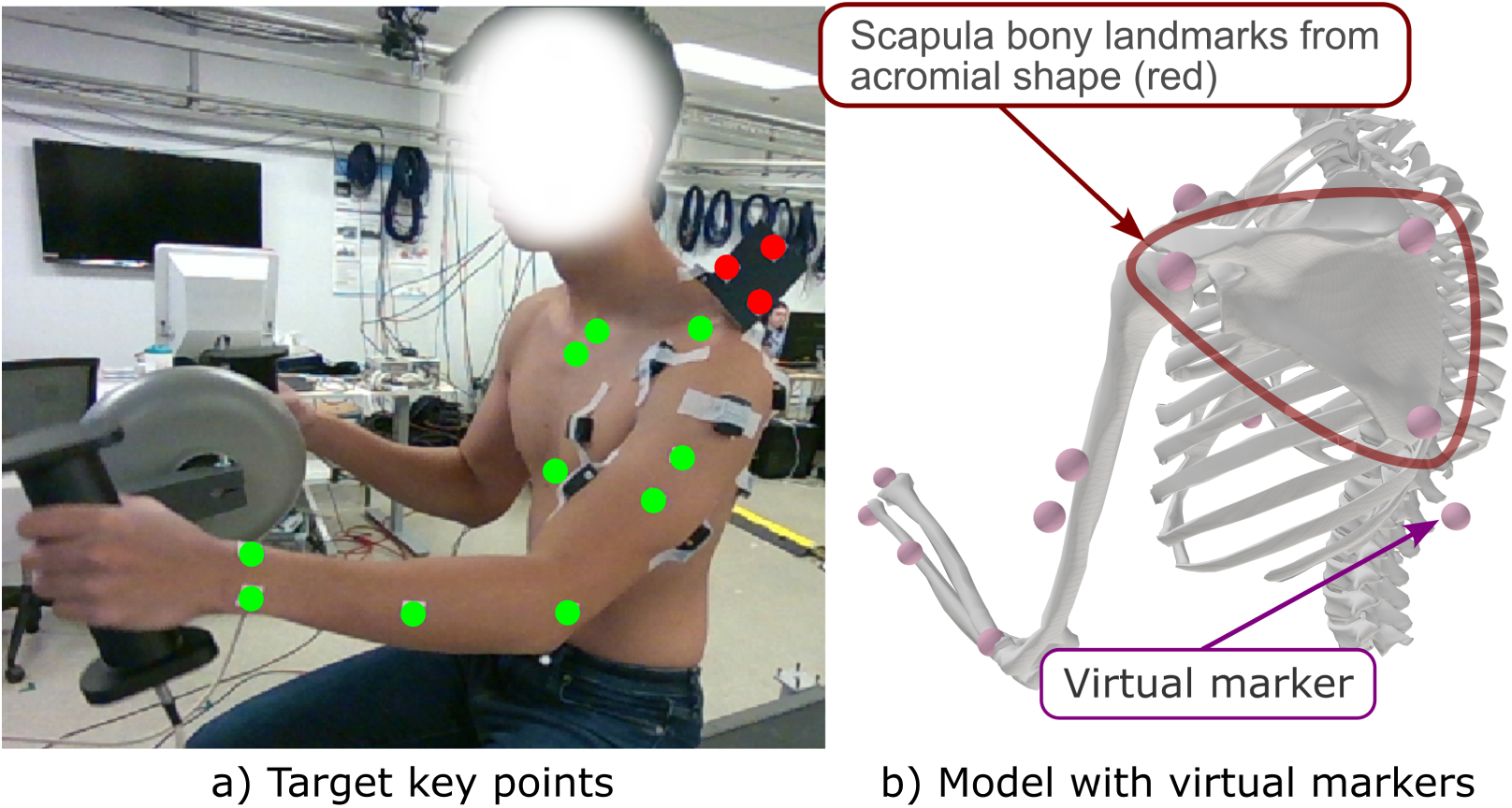
Fourteen Key points extracted from the detection pipeline (left) with the corresponding model (right). The detection included 10 bony landmarks extracted from DeepLabcut on the depth image (green), 3 markers from the acromial shape (red), and the virtual marker. The scapula bony landmarks derived from the acromial shape and the technical marker were incorporated into the model (as shown on b).

### 2.3. Model fine-tuning and validation

The DLC software, providing an interface to pre-trained CNN models such as MobileNet-v2 (Sandler et al., 2018) was used for image key points’ detection. This pre-trained model is relevant to our application as it is compact, can provide low-latency detection, and can be fine-tuned with a limited amount of data (Mathis et al., 2018). In the labeling algorithm of Ceglia et al. (2025), the visibility of the white markers was assessed for each frame to determine the reliability of the detection. Frames in which at least one marker was not properly detected were not considered in the dataset. Ultimately, we decided to keep 500 images per participant (i.e., 4 000 images in total) to reduce the training time and avoid overfitting, this is motivated by the repetitive and constrained aspect of the hand-cycling motion pattern resulting in a limited motion variability. The DLC Python API was used to generate the annotated dataset in the proper format for training. The hyperparameters and data augmentations used for fine-tuning were set to the default ones, except for those presented in Tab. B.4. The fine-tuning was performed using the data of seven participants (3 500 training images), and the validation was made using only the data of the eighth participant (500 validation images), as described in Section 2.5.

### 2.4. Pose estimation

The code used for the pose estimation and results validation is available on Github (Ceglia, 2024). It encompasses the bony landmarks detection from CNN model (Sec. 2.4.1) and, the skeletal-based pose estimation (Sec. 2.4.2).

#### 2.4.1. Bony landmarks detection

Live inference of the fine-tuned Mobilnet-V2 model was used for detecting 13 bony landmarks (Fig. 2) on the depth images through the DLC-Live software (Kane et al., 2020). Since this model does not leverage the time-dependent trajectory of bony landmarks, it may occasionally result in inaccurate location. Additionally, because detection is directly performed on the depth images, a small detection error of a few pixels could result in an incorrect depth value of over one meter (e.g., a bony landmark wrongly located in the background). To address these issues, the bony landmarks detection was integrated into a customized version of our labeling algorithm presented in Ceglia et al. (2025) (Fig. 3). Moreover, since the acromial shape is needed to record personalized scapula kinematics, the white markers placed on it were used during the detection to enhance accuracy. Furthermore, the limited field of view of the RGB-D camera prevents the detection of bony landmarks on the back of the thorax, potentially leading to inaccurate thorax orientation compared to the reference that uses markers on C7 and T10. To address this limitation, a virtual marker is derived from the point cloud provided by the RGB-D camera to enhance thorax orientation estimation. Specifically, 15 three-dimensional points were randomly selected within an area (60×35 pixels) around the sternum bony landmark detected by the CNN model to reconstruct a planar surface. The normal vector to this surface, passing through the detected sternum bony landmarks and oriented posteriorly, was used to define the virtual marker positioned at an empirical distance of 23 cm from the sternum. This virtual marker was incorporated into the skeletal model for each participant (Fig. 2).

**Figure 3:**
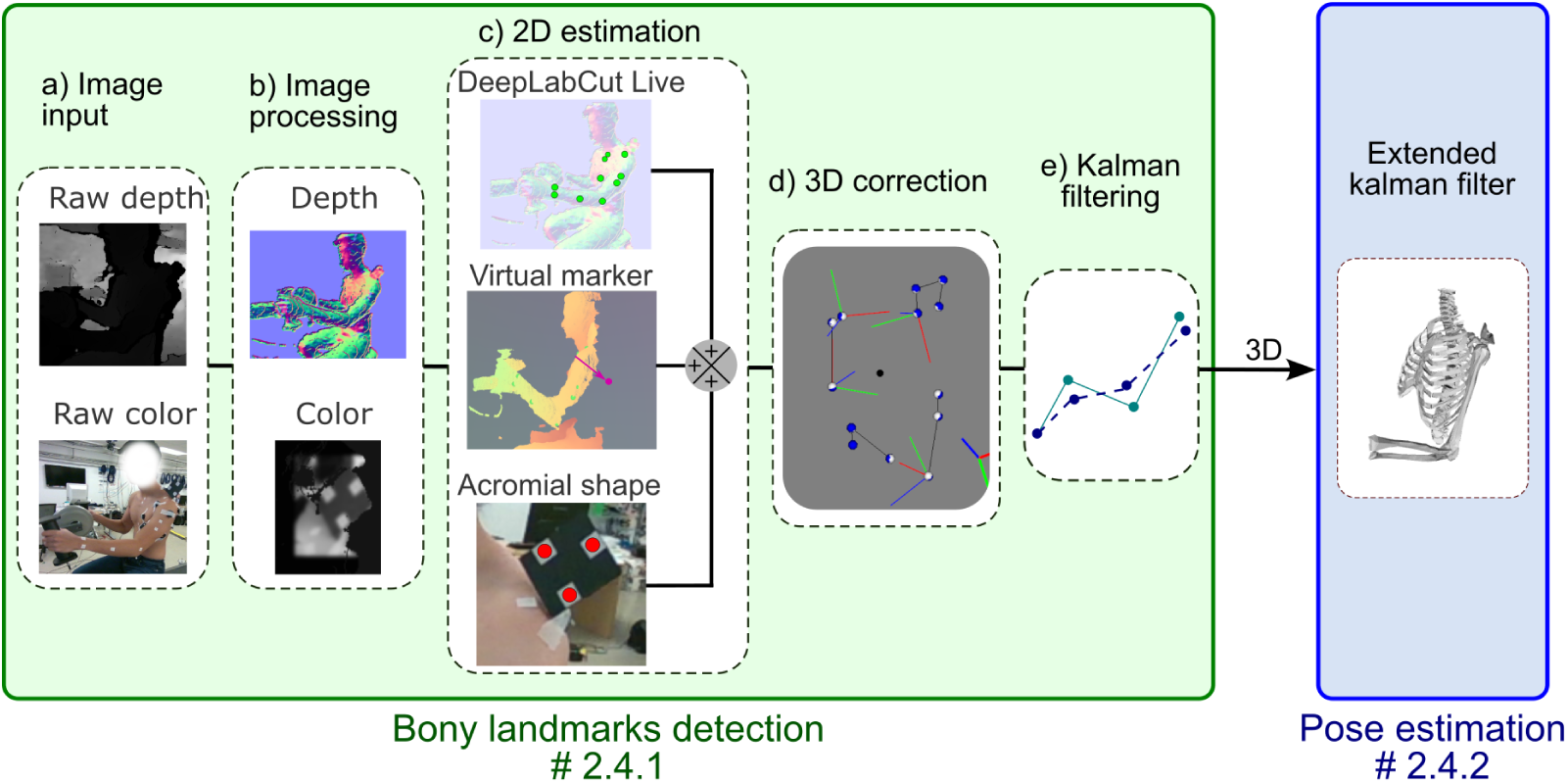
Markers’ detection and pose estimation from the depth and color images.

Finally, our custom detection algorithm consists of five steps (Fig. 3):

a. *Image input*: The raw color and depth images were cropped.
b. *Image processing*: The background was removed on depth and color images using depth threshold. Surface normals (Eq. 2b) were computed from the depth image and fed to the trained DLC model for inference. To facilitate the detection of scapula cluster markers, a contrast-limited adaptive histogram equalization filter was applied to the color image, followed by a blurring process. The settings for color image processing were fine-tuned following the method explained in Ceglia et al. (2025) to detect the white markers on the acromion cluster, the white markers on the skin being ignored.
c. *2D estimation*: The 2D positions of 13 detected bony landmarks (10 from DeepLabCut and 3 from the acromial 3D shape) were converted into meters and expressed within the camera’s 3D coordinate system using the camera’s Python SDK. The default intrinsic matrix provided by the manufacturer was used for this process. Subsequently, the virtual marker position was computed using the point cloud data and the 3D position of the sternum bony landmark.
d. *3D correction*: The 3D markers positions were corrected using a rigid-body model, as described in Ceglia et al. (2025). Once the 3D positions of the markers on the acromion cluster were obtained, they were used to locate the scapula bony landmarks based on measurements made with the device (Appendix A).
e. *Kalman filter*: Finally, as previously reported in Ceglia et al. (2025), the camera operates at a non-conservative 60 Hz frame rate, which can lead to missing frames and therefore missing marker positions. Additionally, the detection algorithm might produce noisy marker trajectories, resulting in unsatisfactory joint velocity estimation. To address these issues, a linear Kalman filter (Kalman, 1960) was employed to predict and fill missing marker positions by estimating the future trajectory point. The filter parameters were determined empirically and applied uniformly across all participants.

These five steps (a-e) provided the 3D positions of 14 bony landmarks (including the virtual marker) (Fig. 2).

#### 2.4.2. Skeletal-based pose estimation

The 14 bony landmarks (Sec. 2.4.1) were used for skeletal-based pose estimation. The upper-limb skeletal model consisted of five segments: thorax, clavicle, scapula, arm, and forearm, and articulated with 16 degrees of freedom (6 at the thorax, 2 at the sternoclavicular joint, 3 at the acromioclavicular joint, 3 at the glenohumeral joint, and elbow flexion and pronation/supination) (Wu et al., 2016). Fourteen virtual markers were placed on the skeletal model to align with the actual bony landmarks (Fig. 2). Initial model scaling was performed using the first five seconds of the hand-cycling motion, leveraging the detected bony landmarks (Sec. 2.4.1) and the Opensim software (Delp et al., 2007). This scaling step ensured that the model was accurately adapted to each participant’s specific anatomy. Once the scaling was completed, the bony landmarks positions were input into an extended Kalman filter, implemented through the Biorbd software (Michaud and Begon, 2021), to estimate joint angles and angular velocities.

### 2.5. Data analysis

An 8-fold cross-validation was performed, where eight models were fine-tuned. Each model was trained using the images from seven participants and validated on the images of the remaining participant, ensuring that every image in the dataset was used once for validation. Bony landmarks detection accuracy was initially assessed in 2D by computing the root-mean-square errors (RMSE) of the detections made by each model relative to color image-based annotations. This analysis evaluated the impact of training dataset variations on the method’s accuracy. Subsequently, RMSE of the 3D positions estimated by our method compared to those obtained from the Vicon system was computed. Bony landmarks extracted from DLC were projected into the Vicon coordinate system using the rototranslation matrix described in Ceglia et al. (2025) for each motion. Synchronization of the systems was achieved using a trigger signal processed with the *biosiglive* library (Ceglia et al., 2023). To align temporal resolution, DLC outputs were interpolated to match Vicon’s 120 Hz frame rate. Root-mean-square differences (RMSD) for joint angles and velocities were also computed relative to the reference system. Agreement between the two systems was further assessed using Bland–Altman analysis (Bland and Altman, 2007), focusing on the *bias* and *limits of agreement* (LoA).

Performance metrics including the latency introduced by marker detection and pose estimation were also evaluated. All tracking algorithms and biomechanical analyses were executed on a workstation equipped with an AMD Ryzen 9 5950X 16-Core 3.40 GHz processor and an NVIDIA GeForce RTX 3060 GPU, running Ubuntu 22.04 LTS.

## 3. Results

The RMSE of the bony landmarks location for each model remained consistent across all subjects (Tab. 1), regardless of sex, with values ranging from 3.83±1.66 to 5.42±1.62 pixels. These results indicate a high level of stability in the model with respect to the training data. The overall error across all models was 4.43±1.35 pixels, less than 1% of the image’s diagonal.

**Table 1:**
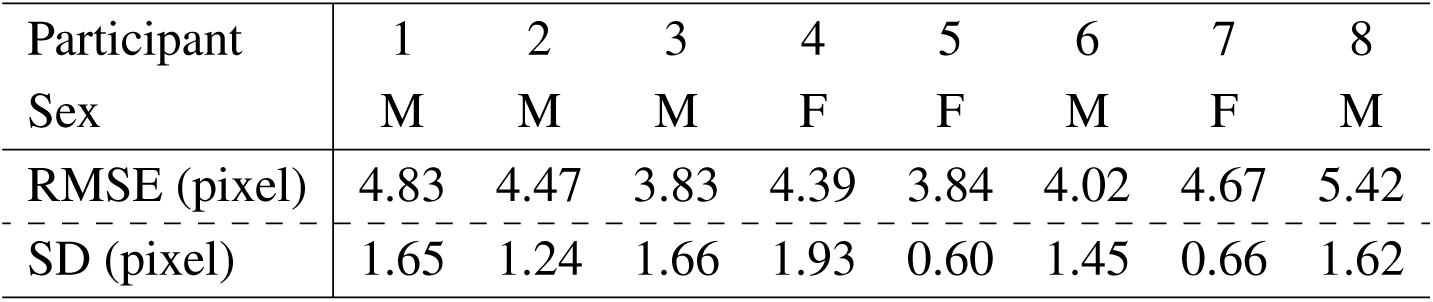
Root-mean-square error (RMSE) and standard deviation (SD) of the detection performed by each model compared to the annotated images.

| Participant | 1 | 2 | 3 | 4 | 5 | 6 | 7 | 8 |
| --- | --- | --- | --- | --- | --- | --- | --- | --- |
| Sex | M | M | M | F | F | M | F | M |
| RMSE (pixel) | 4.83 | 4.47 | 3.83 | 4.39 | 3.84 | 4.02 | 4.67 | 5.42 |
| SD (pixel) | 1.65 | 1.24 | 1.66 | 1.93 | 0.60 | 1.45 | 0.66 | 1.62 |

The average error in 3D trajectories of the bony landmarks compared to the Vicon-based motion capture system, was 5.46±3.76 mm across all participants (Tab. 2). Bland-Altman analysis demonstrated strong agreement, with a bias of −0.80 mm and limits of agreement ranging from −8.52 mm to 6.92 mm. The propagation error was minimal, with RMSD values for joint angles and velocities averaging 3.81±1.67° and 2.81±2.16°/s, respectively. Bland-Altman analysis indicated near-zero bias for both joint angles and velocities, with relatively narrow limits of agreement: (−10.70°, 10.34°) for joint angles and (−8.74°/s, 8.44°/s) for velocities. The largest differences in joint angle estimation were observed for the humerus plane of elevation (10.67°), humerus axial rotation (9.89°), and forearm pronation/supination (7.04°) (Fig. 4). For other joints, the mean differences remained below 6° (Fig. 4).

**Figure 4:**
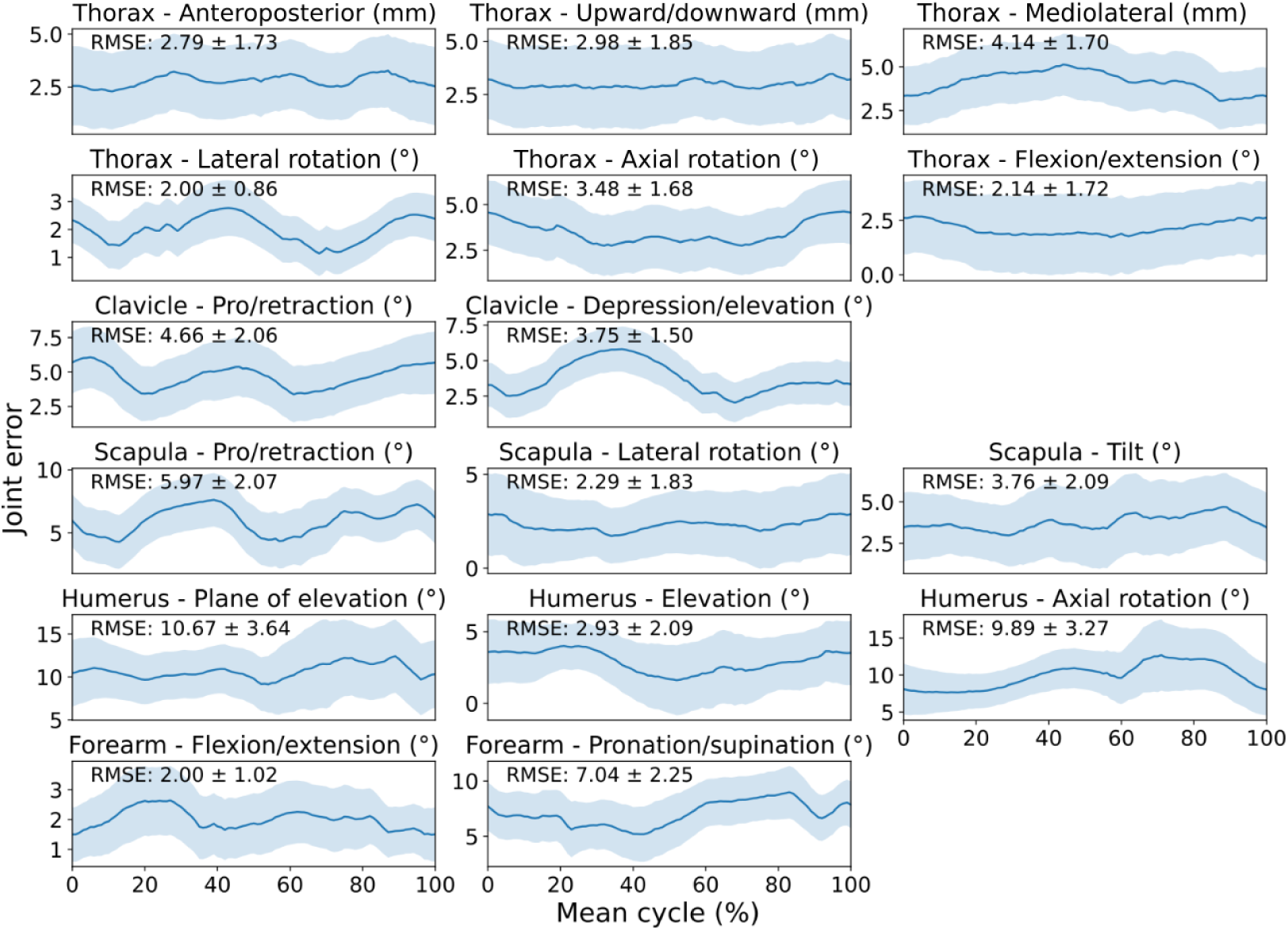
Error in joint angle estimation using the proposed method compared to the marker-based system. The error is depicted as the mean (solid line) ± one standard deviation (shaded area) across all cycles, participants, and trials for each degree of freedom. The root-mean-square errors (RMSE)±one standard deviation is reported for each degree of freedom.

**Table 2:** Root Mean Square Deviation (RMSD) and one standard deviation (SD), along with Bland-Altman limit of agreement (LOA), LOA 95% confidence interval (CI), and bias, of the 3D marker trajectories, joint angles and velocities.

|  | RMSD SD |  | LOA |  | CI | Bias |
| --- | --- | --- | --- | --- | --- | --- |
|  |  |  | Low | High |  |  |
| Markers (mm) | 5.46 | 3.76 | -8.52 | 6.92 | 1.31 | -0.80 |
| Joint angles (°) | 3.81 | 1.67 | -10.70 | 10.34 | 1.26 | -0.18 |
| Joint velocities (°/s) | 2.81 | 2.16 | -8.74 | 8.44 | 0.84 | -0.15 |

A key aspect of this work is the algorithm’s performance in delivering real-time pose estimation. The detection of bony landmarks and computation of technical markers from the surface (Sec. 2.4.1) were achieved with a latency of 18.64±2.57 ms (*i.e.*, **53** Hz) (Tab. 3) per image. The inverse kinematics computation required approximately 1.23 ± 0.06 ms per frame. Overall, the pose estimation algorithm generated 3D marker trajectories and joint kinematics with a total latency of 19.88 ± 2.63 ms (*i.e.*, **50** Hz) (Tab. 3) per frame, slightly below the 60 frames per second provided by the camera.

**Table 3:** Mean and standard deviation (SD) of the latency introduced by the algorithm for the bony landmarks’ detection, computing the virtual marker and inverse kinematics for one image.

|  | Mean (ms) | SD (ms) |
| --- | --- | --- |
| Bony landmarks' extraction | 17.53 | 2.21 |
| Virtual marker computation | 1.11 | 0.36 |
| Pose estimation | 1.23 | 0.06 |
| Total | **19.88** | **2.63** |

## 4. Discussion

This study aimed to propose a real-time markerless method for refined upper-limb kinematics using a single RGB-D camera by combining automatic detection of bony landmarks with a custom acromial 3D shape. The proposed approach was evaluated against a reference optoelectonic marker-based system to assess its accuracy in bony landmark location and error propagation in kinematic outcomes, including joint angles and velocities. The method showed accurate bony landmark detection with minimal propagated error in real-time (50 Hz), making it a realistic and cost-effective alternative to traditional marker-based motion capture systems. Additionally, the inclusion of a custom 3D shape enabled scapula integration into the kinematic model, highlighting the potential for clinical applications. All software and hardware developments made to achieve our goal, including the detection pipeline and the acromial shape, are freely available to broaden their use and facilitate practical applications.

### 4.1. Bony landmark detection

The method provided reliable detection of bony landmarks, with an average error of less than 6 mm compared to the marker-based system. This error level is comparable to the effects of soft tissue artifacts (up to 18 mm (Ancillao et al., 2021)) and falls within the range of marker placement errors (up to 9 mm (Salvia et al., 2009)). This minimal error was achieved through the fine-tuning of a CNN model (Sandler et al., 2018), supported by the high-quality annotations of the training images. Bony landmarks were manually identified and marked on the participant and then accurately detected in the RGB images to annotate the depth images (Ceglia et al., 2025). This depth annotation method, using one RGB-D sensor and some white markers, can easily be used to create datasets for depth-based keypoints detection models. Traditionally, such annotations are performed either by manually identifying points in images, that lack precision, or by using reflective markers removed via machine learning techniques, that can alter training images. In contrast, our method simplifies the annotation process while maintaining accuracy, improving its practicality. Only the acromion cluster needs to be attached, no markers are required to be placed on the body for kinematics estimation. The custom algorithm for bony landmark extraction (Sec. 2.4.1) further reduced error by making it possible to use color markers on the acromion cluster (via RGB image) and integrating a 3D rigid-body skeleton, ensuring reliable marker positions over time. Additionally, since only the depth image is used for detecting bony landmarks (excluding the acromion cluster), the method is not sensitive to variations in skin color and brightness.

### 4.2. Error propagation

The Bland-Altman analysis revealed a low bias for joint angles (−0.18°), with an average error of 3.81±1.67° for joint angle estimation. This level of error is significantly lower than the 30° reported for upper-limb motion (Henschke et al., 2022) and comparable to the 10° reported for gait using IMU methods (Shuai et al., 2022). Additionally, our results outperform other computer vision methods, which have error margins reaching up to 23° for upper-limb joint angles (Lahkar et al., 2022). The use of a custom-made acromial 3D shape enabled scapula motion estimation, with errors ranging from 2° to 6°. This capability allows for a complete kinematic analysis of all upper-limb joints, in contrast to existing methods that reduce the shoulder complex to a single humerothoracic joint (Cai et al., 2019; Lahkar et al., 2022; Boldo et al., 2024). Such comprehensive kinematics make our method particularly suitable for clinical evaluations of shoulder disorders. While it is challenging to determine if the reported errors are universally low enough, they might be sufficient for detecting clinically relevant changes, such as the minimum detectable change for scapular winging(∼5° (Larsen et al., 2020)). However, the method requires the placement of an additional cluster on the acromion. Although commonly used for scapula motion capture, this requirement may not always be practical in certain contexts. Finally, by leveraging the RGB-D camera’s point cloud, an additional virtual marker was computed, which reduced the error in thorax 3D positioning. This highlights the potential of the RGB-D camera for enhancing accuracy.

### 4.3. Latency

The algorithm operated with a total latency of 19.88±2.63 ms, which is slightly lower than the 60 Hz camera frame rate (16 ms latency). This performance was achieved using the MobileNetV2 CNN model, chosen for its efficiency in real-time inference. Nevertheless, the model was not optimized for its fastest configuration. Further improvements could be achieved by using inference optimization methods such as those provided by TensorFlow Lite (David et al., 2021), which, while untested here, could significantly enhance inference performance. Additional latency reductions could be achieved by exploiting the multiprocessing capabilities of CPUs. For example, processing depth and color images in separate threads or handling scapula cluster marker detection and DeepLabCut inference in parallel could improve overall processing speed. These optimizations may help the system exceed the camera frame rate and further enhance its real-time performance. To address the occasional instability in frame rate, a Kalman filter was implemented to predict and fill gaps in the data, effectively mitigating the impact of frame drops. Despite these areas for improvement, the current total latency of approximately 20 ms is well below the recommended threshold for effective visual biofeedback, which ranges between 30 and 85 ms (Kaaresoja et al., 2014). This ensures the method remains suitable for real-time applications.

### 4.4. Limitations

This study has several limitations that should be acknowledged. The dataset comprised a small number of participants, limiting variability and increasing the potential for overfitting, even though a pre-trained model was fine-tuned rather than trained from scratch. To mitigate this, a cross-validation method was employed to evaluate the model on previously unseen data. Nevertheless, the low RMSE reported in Table 2 resulting from models trained in low-data regime are highly encouraging. This can be explained by the low variability in the somehow constrained hand-cycling motion pattern. However, only healthy participants were recruited, despite the technology being intended for clinical populations. Applying the model to pathological motion without additional training on relevant data may yield inaccurate results.

Participants in this study were either shirtless or wearing sports bras, so the algorithm’s performance with regular clothing remains uncertain. Additional experiments are required to evaluate the performance of our method on clothed participants. Furthermore, data collection occurred in a controlled environment with consistent lighting and minimal background distractions. Although the algorithm systematically removes backgrounds and the stereo vision camera is designed to be robust to lighting and environmental variations, testing in more diverse conditions will be necessary to confirm robustness.

The algorithm requires a computer equipped with a GPU to achieve real-time performance. Although these GPUs are now relatively affordable (approximately $500), their use could still complicate the application in real-world scenarios or on embedded devices. While it can run accurately on a CPU, the reduced operational rate (∼10–20 Hz) may not meet the demands of certain applications. Lastly, the method was tested only on repetitive motions with minimal occlusions, leaving its robustness in more complex movements with significant occlusions unassessed. Future work should address these aspects to enhance the method’s applicability in clinical contexts.

## 5. Conclusion

We proposed a novel method for refined upper limb kinematics estimation using a single RGB-D camera combined with an acromial cluster. Our approach enabled real-time bony landmarks detection and inverse kinematics, which show a strong agreement with a standard marker-based method. This highlights the potential of using an accurate, low-cost, user-friendly system to assess the upper limb movement including the shoulder complex in clinical contexts. A feasibility study with patients in a more representative context will follow.

## Appendix A. Description and evaluation of the custom acromial 3D-shape

### Appendix A.1. Device description

The acromial shape, designed in SolidWorks (Dassault Systèmes, France), comprises two components (Fig. A.5): 1) a **location system** (acromial cluster) and 2) a **pointing system**. Both components are designed for 3D printing and are compatible with either the right or left scapula. The device used in this study was a prototype, and all parts are freely available in STL format, along with detailed assembly and usage instructions (https://github.com/aceglia/scapula_cluster/tree/main).

**Figure 5:**
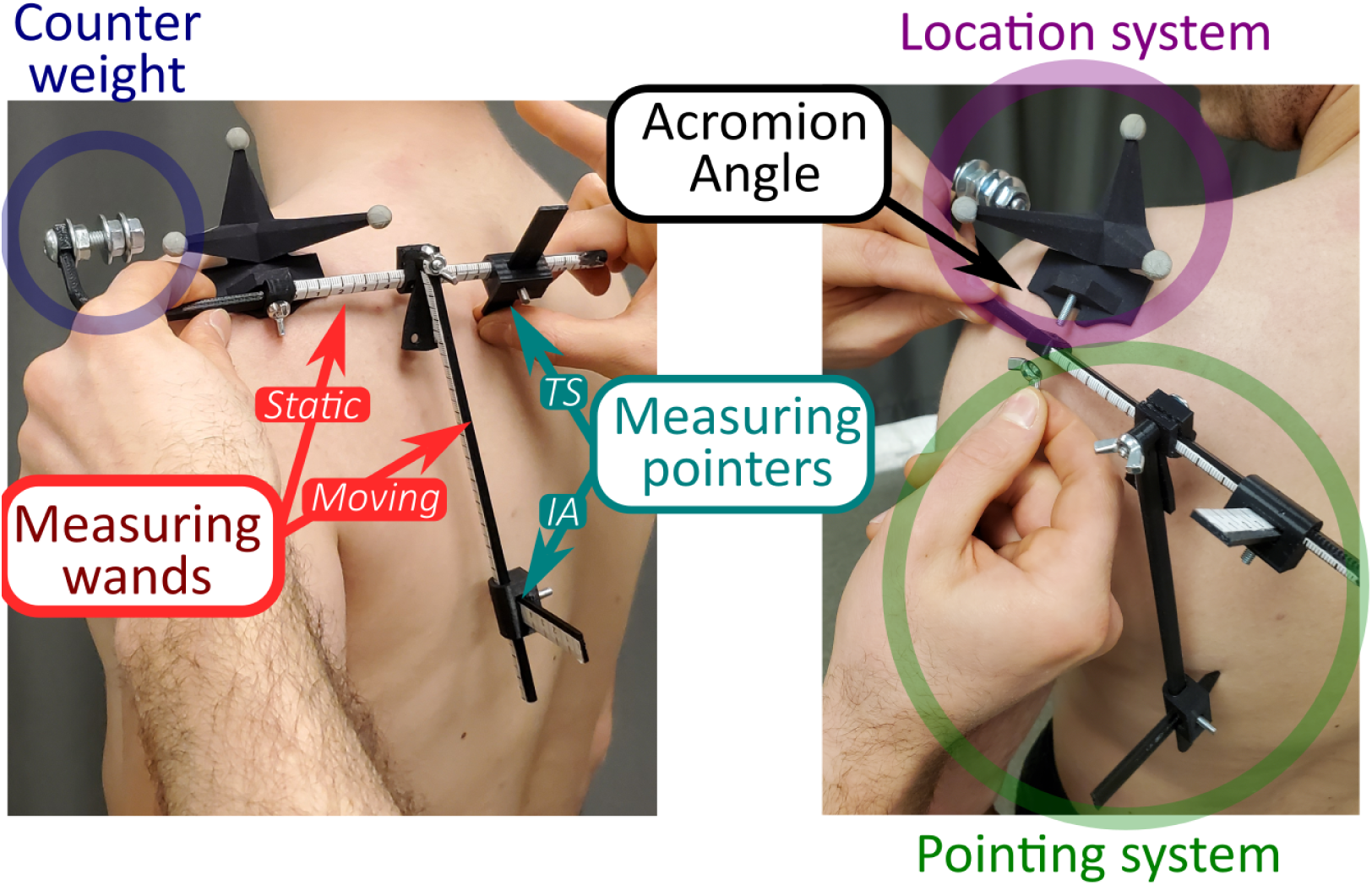
Example of the device being placed (left) on a participant, and the pointing system and the counterweight being removed (right).

The **location system** is intended to capture the scapular orientation. It consists of a customizable 3D shape integrated with a tracking system (Fig. A.5). This 3D shape is positioned at the most lateral, supero-posterior part of the scapular spine, starting medially at the acromial angle (AA). For evaluation, a set of reflective markers was used as the tracking system, though the study employed a custom-designed 3D shape tailored for use with a single RGB-D camera (Appendix A.3). The chosen tracking system enables the definition of a local coordinate system centered on AA to record the scapular orientation during movement.

The **pointing system** is designed to perform the anatomical calibration of the location system. It consists of two wands and two pointers equipped with measuring tapes, and a counterweight to ensure device balance (Fig. A.5). During the calibration process, the pointing system is securely attached to the location system via a screw connection. After proper adjustments, the pointers are placed on the skin over the trigonum spinae (TS) and inferior angle (IA). This setup allows for the calculation of the landmark positions within the local coordinate system using matrix transformations.

To position and calibrate the device while the participant is in the anatomical position, the following steps should be followed:

1. Secure the device onto the most lateral, supero-posterior part of the scapular spine. Ensure that AA is in contact with the latero-posterior corner of the location system base and align the static wand with the scapular spine.
2. Position the TS pointer over TS. Ensure that the tip gently touches the skin’s surface without pressing too deeply. Tighten the screw to secure the pointer in place.
3. Align the moving wand with IA and tighten the screw to fix its position.
4. Position the IA pointer over IA in the same manner as the TS pointer, and secure the position.
5. Remove the pointing system by loosening the screw on the base.

Once the location system is secured to the skin and the pointing system has been removed, data collection can begin. At a convenient moment, the user should retrieve five measurements from the measuring tapes of the pointing system: the two distances along the wands, the two depths of the pointers, and the distance from the moving wand to the base. These values are then input into an open-source Python script containing preloaded transformation matrices. The script computes the positions of the anatomical landmarks (AA, TS, and IA) within the coordinate system of the motion capture system. The calculations are based on the chosen tracking system, with the relationship between the 3D shape orientation and the base coordinate system used to determine the global positions of the landmarks. For more information on the Python code and the transformation matrix definitions, readers can consult the documentation and code available on GitHub (https://github.com/aceglia/scapula_cluster).

### Appendix A.2. Evaluation against scapula locator

To assess the accuracy of our acromial cluster in retrieving the scapular orientation, the criterion validity against a scapula locator (van Andel et al., 2009; Brochard et al., 2011; Bet-Or et al., 2017) was estimated. Eight healthy adults (3 females; age: 25.9±12.2 years; height: 175.8±8.5 cm; mass: 70.2±8.4 kg) participated in the study. This experiment was approved by the ethics committee of the Université de Montréal (#2023-4743). The acromial cluster was first positioned on the left shoulder and calibrated following the procedure previously outlined. The arm position was then adjusted using a goniometer at elevations of 0°, 45°, 90°, and 120° in both the sagittal and frontal planes. The maximum arm elevation angle was limited to 120° since previous studies have shown that the acromion cluster’s orientation estimation diverges from invasive methods at arm elevation angles exceeding 120° (Karduna et al., 2001). For each static position, the scapular landmarks (AA, TS, and IA) were manually palpated, and a scapula locator was used to provide reference positions of these landmarks. Measurements were taken twice, with a 30-second rest between trials. A 15-camera Vicon system (Oxford, UK) was used to record at 120 Hz the trajectories of the markers placed on the acromion cluster and the scapula locator. The scapular orientation was computed according to the recommendations of the International Society of Biomechanics (ISB) (Wu et al., 2005) and was reported in terms of retro/protraction, lateral rotation, and anterior/posterior tilt. To quantify the accuracy, the root-mean-square error (RMSE) was calculated to compare the scapular orientation estimated by the acromion cluster and the scapula locator across all participants and trials for each Euler angle.

The overall average error across all axes was 3.08±4.12°, with the highest error observed for the rotation angle (3.17±5.41°), followed by the retraction angle (3.10±2.77°) and the tilt angle (2.97±4.18°). The deviation angles between the scapular orientations estimated by the two systems were also analyzed to provide insight into the direction of the difference (Fig. A.6). This analysis revealed that, for all axes, the differences were centered close to zero, although a slight trend toward higher protraction estimates by the acromial cluster was observed (Fig. A.6).

**Figure 6:**
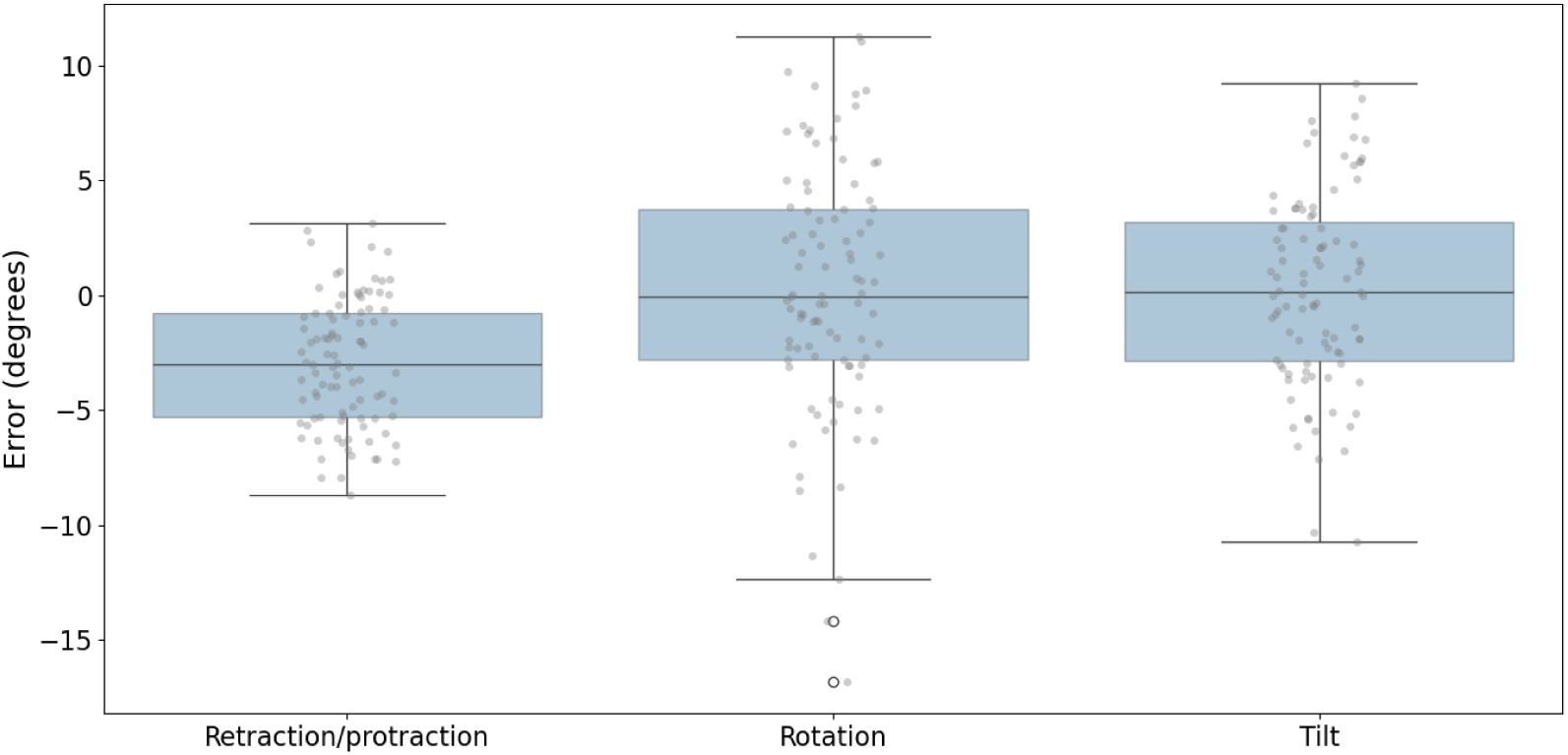
Box plot displaying the interquartile range (25th and 75th percentiles) of the error between the cluster and the scapula locator for each Euler angle across all positions and participants.

Our evaluation was conducted on a limited number of participants and did not include a reliability assessment. However, we found that our acromial cluster provides scapular orientation estimates comparable to those obtained with a scapula locator (van Andel et al., 2009; Brochard et al., 2011; Bet-Or et al., 2017). Furthermore, the 25th and 75th percentiles for all Euler angles fall between −6° and 5°, remaining within an acceptable margin of error for most applications. In addition, the device design is freely available and can be 3D-printed at a cost of less than $150. While further tests and refinements are necessary, this work paves the way for the potential use of a standardized acromial cluster in both marker-based and markerless motion capture systems.

### Appendix A.3. Custom base

The acromial cluster was designed to be customizable, allowing it to be modified and adapted for specific applications. In our primary study, it consisted of a base (common to all custom location systems) and a screen visible to the RGB-D camera from a frontal view (Fig. 3). The coordinate system derived from the three white markers on the screen was linked to the local coordinate system of the acromial cluster. A transformation matrix allowed for the conversion of the anatomical landmark positions from the local coordinate system of the acromial cluster to the global coordinate system of the motion capture system. For any newly designed custom 3D shape, this transformation matrix must be determined and integrated into the Python script to ensure accurate computation of anatomical landmark positions in the global coordinate system.

## Appendix B. Settings of DeeplabCut model fine-tuning

The default values provided by DeeplabCut for fine-tuning the CNN model were mostly kept except for the ones presented in the table below (Tab. B.4). Specifically, we retained augmentations related to the orientation and scaling of the image and removed all augmentations related to contrast and image quality.

**Table B.4:** Default and custom parameters for model fine-tuning and data augmentation with DLC, with justifications when relevant.

|  | Name | Default value | Custom value | Justification |
| --- | --- | --- | --- | --- |
| Image augmentation | Convolution | Emboss | None | Image from surface normals |
|  | Contrast | [Clahe, Histeq] | None | Keep image original color |
|  | Crop sampling | Hybrid | Density | All features in the same image space |
|  | Elastic transform | True | False | Keep image proportion |
|  | Blur | True | False | Use downsampling to reduce quality |
|  | Scale jitter | [0.5, 1.25] | [0.8, 1.1] | Image cropped around the participant |
|  | Global scale | 0.8 | 1 | Downsampling made manually on dataset |
| Training | Net type | Resnet-50 | MobilnetV2-0.35 | Efficiency |
|  | Max iteration | None | 600 000 | Reach the plateau |
|  | Key points | None | 13 | - |

## Declaration of generative AI and AI-assisted technologies in the writing process

During the preparation of this work the authors used Grammarly, ChatGPT, and LanguageTool in order to correct spelling and grammatical errors and improve text clarity and flow. After using these tools, the authors reviewed and edited the content as needed, and take full responsibility for the content of the publication.

